# Featural representation and internal noise underlie the eccentricity effect in contrast sensitivity

**DOI:** 10.1101/2023.04.04.535413

**Authors:** Shutian Xue, Antonio Fernández, Marisa Carrasco

## Abstract

Human visual performance for basic visual dimensions (e.g., contrast sensitivity and acuity) peaks at the fovea and decreases with eccentricity. The eccentricity effect is related to the larger surface area of the visual cortex corresponding to the fovea, but it is unknown if differential feature tuning contributes to this eccentricity effect. Here, we investigated two system-level computations underlying the eccentricity effect: featural representation (tuning) and internal noise. Observers (both sexes) detected a Gabor embedded in filtered white noise which appeared at the fovea or one of four perifoveal locations. We used psychophysical reverse correlation to estimate the weights assigned by the visual system to a range of orientations and spatial frequencies (SFs) in noisy stimuli, which are conventionally interpreted as perceptual sensitivity to the corresponding features. We found higher sensitivity to task-relevant orientations and SFs at the fovea than the perifovea, and no difference in selectivity for either orientation or SF. Concurrently, we measured response consistency using a double-pass method, which allowed us to infer the level of internal noise by implementing a noisy observer model. We found lower internal noise at the fovea than perifovea. Finally, individual variability in contrast sensitivity correlated with sensitivity to and selectivity for task-relevant features as well as with internal noise. Moreover, the behavioral eccentricity effect mainly reflects the foveal advantage in orientation sensitivity compared to other computations. These findings suggest that the eccentricity effect stems from a better representation of task-relevant features and lower internal noise at the fovea than at the perifovea.

**Significance:** Performance in many visual tasks worsens with eccentricity. Many studies attribute this eccentricity effect to retinal and cortical factors, like higher cone density and a larger cortical surface area representing the foveal than peripheral locations. We investigated whether system-level computations for task-relevant visual features also underlie this eccentricity effect. Measuring contrast sensitivity in visual noise, we showed that the fovea better represents task-relevant orientation and spatial frequency and has lower internal noise than the perifovea, and that individual variability in these two computations correlates with that in performance. These findings reveal that both representations of these basic visual features and internal noise underlie the difference in performance with eccentricity.

## Introduction

Visual performance worsens with eccentricity for many visual dimensions, including contrast sensitivity (Cannon, 1985; Baldwin et al., 2012; Jigo et al., 2023) and acuity (Strasburger et al., 2011; Anton-Erxleben and Carrasco, 2013). The foveal advantage relates to higher cone density (Polyak, 1941), smaller retinal receptive fields (Enroth-Cugell and Robson, 1966), and larger V1 area processing the stimuli (Virsu and Rovamo, 1979; Duncan and Boynton, 2003; Benson et al., 2021; Himmelberg et al., 2021). However, it is unknown whether system-level representations and computations for task-relevant features differ across eccentricities, and whether they underlie the eccentricity effect in contrast sensitivity. Thus, we investigated three questions:

1. Does the representation of orientation and SF differ between fovea and perifovea? Representation is considered the externalization of the internal neural processes elicited by perceptual and cognitive processes. Functionally, it serves as a template for observers to perform a given task (Ahumada, 2002; Gold et al., 2004; Nagai et al., 2008) and its efficiency depends on resemblance to the signal (Burgess et al., 1981; Levi and Klein, 2002) and the ideal observer template (Abbey and Eckstein, 2009). This representation indicates the system’s tuning properties–the sensitivity to and selectivity for features–and approximate electrophysiological sensory tuning properties (Neri and Levi, 2006). Representation modulations reflecting differential system-level computations underlie performance changes due to expectation (Wyart et al., 2012) and covert spatial (Fernández et al., 2019, 2022), feature-based (Paltoglou and Neri, 2012) and presaccadic (Li et al., 2016; Ohl et al., 2017) attention. Here, we investigated the representation of two fundamental visual features– orientation and spatial frequency–which are jointly encoded (Blakemore and Campbell, 1969; De Valois et al., 1982a; Moraglia, 1989) and processed by parallel channels tuned for orientation and SF (De Valois and De Valois, 1988; Graham, 1989). We used psychophysical reverse correlation (Abbey and Eckstein, 2002; Ahumada, 2002; Fernández et al., 2019, 2022; Wyart et al., 2012) in conjunction with a detection task to simultaneously derive the representation of both orientation and SF.
2. Does internal noise differ between fovea and perifovea? Our visual system is perturbed by internal noise, as reflected in contrast (Pelli, 1985; Legge et al., 1987; Ahumada, 2002), orientation (Heeley et al., 1997; Lu and Dosher, 1998) and motion (Ling et al., 2009) tasks. Most studies estimate the level of internal noise only at the fovea (Burgess and Colborne, 1988; Lu and Dosher, 1999) or periphery (Dosher and Lu, 2000; Eckstein et al., 2004; Mareschal et al., 2008b). When eccentricity increases, internal noise has been reported to increase (motion: Mareschal et al., 2008a) or remain constant (motion: Falkenberg and Bex, 2007; face: Mäkelä et al., 2001). Whether internal noise increases with eccentricity for tasks mediated by basic visual features (e.g., orientation and SF) is unknown. We investigated whether internal noise limits the processing of basic visual features–orientation and SF–between fovea and perifovea. First, we inferred the internal noise level from response consistency, assessed with a double-pass method via multiple presentations of identical noisy stimuli (Burgess and Colborne, 1988). Second, we estimated the internal noise level by implementing a noisy observer model, which predicts trial-wise responses based on the measured response consistency.
3. Do differential computations mediate behavioral differences between fovea and perifovea? Individual differences in contrast sensitivity correlate with surface area in early occipital areas (Himmelberg et al., 2022), and with internal noise in humans (McAnany and Park, 2018) and macaques (Kiorpes et al., 1999). We tested whether the eccentricity effect can be accounted for by the location difference in computations by assessing the correlation between the individual variability in computation and behavior. In sum, we investigated system-level computations underlying the eccentricity effect in contrast sensitivity by: (1) comparing the representations of orientation and SF and the level of internal noise between fovea and perifovea, and (2) testing the hypothesis that the eccentricity effect in contrast sensitivity reflects discrepancies in featural representation and internal noise between fovea and perifovea.

## Materials and Methods

### Participants

Twelve observers (8 females, age: 23-28, mean=26.25, SD=2.36) participated in the study. Ten were experienced psychophysical observers and all but one (author, SX) were naïve as to the purpose of this study. All observers provided written informed consent and had normal or corrected-to-normal vision. All experimental procedures agreed with the Helsinki declaration and were approved by the University Committee on Activities Involving Human Subjects at New York University.

### Apparatus

All stimuli were generated and presented using MATLAB (MathWorks, Natick, MA) and the Psychophysics Toolbox (Kleiner et al., 2007) on a gamma-linearized CRT monitor (1280×960 screen resolution; 100 Hz; 33 cd/m2 background luminance). Observers viewed the display at 57 cm with their heads stabilized by a chin rest. An eye-tracker system (EyeLink 1000) was in front of the observer to track eye position.

### Experimental Design

#### Stimuli

Figure 1 illustrates one trial. A black fixation cross (arm length=0.3º) was presented at the center of the screen before the stimulus onset. Five black placeholders, each composed of four corners (line length=0.75°, delimiting a virtual 4°x4° square) were simultaneously presented at the fovea and four isoeccentric perifoveal locations at 6º eccentricity: two along the horizontal meridian and two along the vertical meridian. Placeholders indicate the location of the target and four distractors with 100% validity to minimize spatial uncertainty because it increases with eccentricity (Hess and Hayes, 1994; Michel and Geisler, 2011). Observers were instructed to report whether they detected a Gabor in the target. In half of the trials, the target consisted of a noise patch (20% RMS contrast, containing SFs within 1-4 cpd with uniform power, width=3º). Note that there was no energy above 4 cpd, but there was some energy below 1 cpd and observers might have used this information; however, the power between 1-4 cpd was ∼96% of the power spectrum and equivalent across locations. And in the other half of the trials, the target contained a horizontally oriented Gabor embedded in the noise patch. The Gabor was generated by modulating a 2-cpd sinewave delimited by a Gaussian envelope (SD of the Gaussian envelope=0.8; width of the noise patch=3º). The phase of the Gabor was random on each trial to avoid adaptation. The root-mean-square (RMS) contrast of the Gabor was determined independently at each location for each observer through a titration procedure to yield 70% accuracy. All four distractors were noise patches independently generated at each location on each trial.

**Figure 1.**
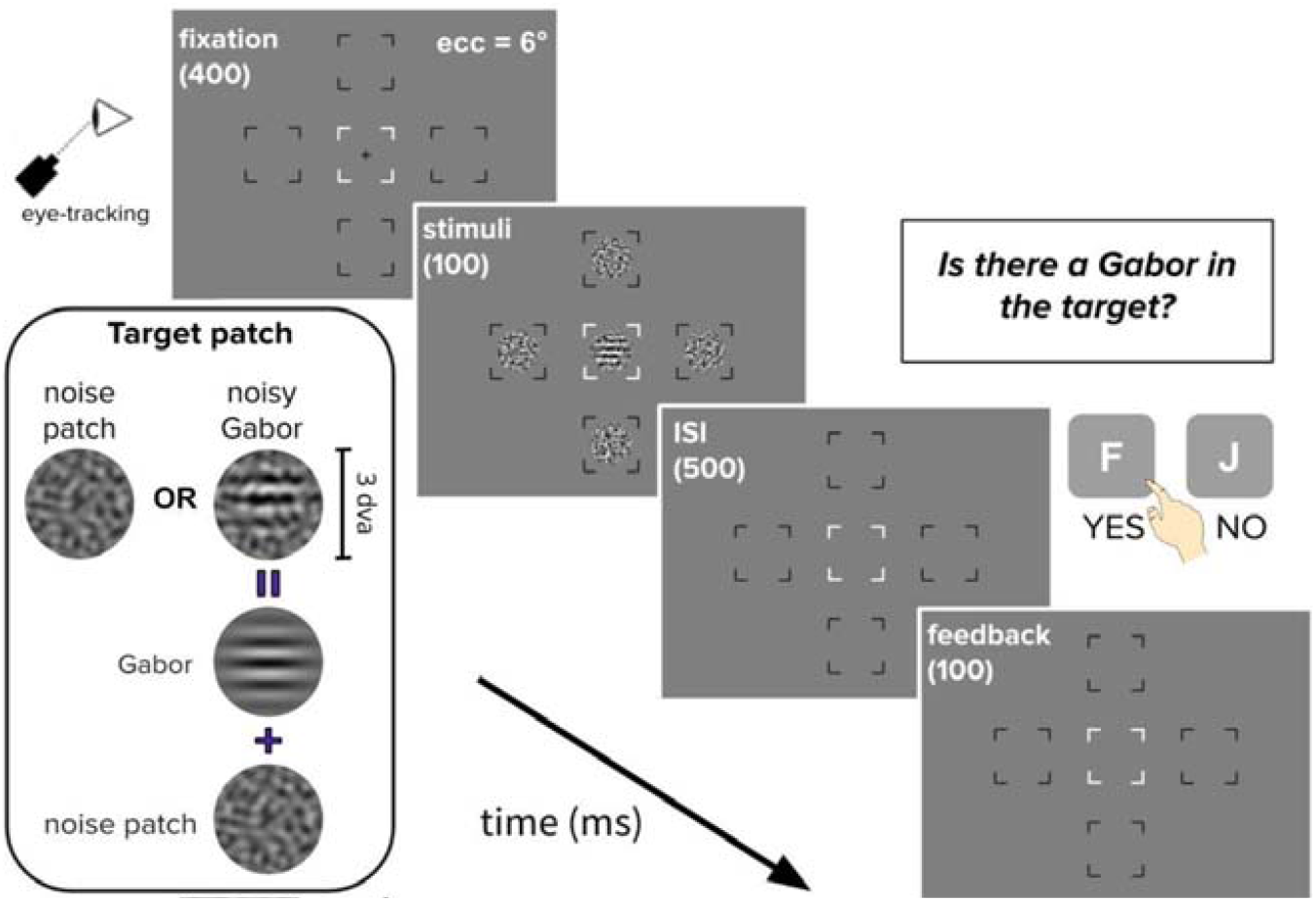
Trial sequence and stimuli. In half of the trials, the target patch (indicated by the white placeholder) consisted of a horizontal Gabor (SF=2 cpd) embedded in filtered-white noise; in the other half, the stimulus was just independently generated noise. Observers performed a yes/no detection task by reporting whether they observed a Gabor in the target. Auditory feedback was provided after the response was given.

#### Procedure

Each trial began with a fixation cross at the center of the screen. Then, five placeholders were presented simultaneously for 400 ms followed by the display of target and distractors for 100 ms. Target location was blocked and observers were informed of the target location at the start of the block to eliminate spatial uncertainty. Placeholders were on the screen until observers made a response. Accuracy was emphasized and there was no time limit to respond. Auditory feedback was given after each response: A high-tone beep indicated correct and a low-tone beep indicated incorrect responses. At the end of each block, accuracy was displayed on the screen.

To infer the amount of internal noise in the system, we measured response consistency by employing the double-pass method (Burgess and Colborne, 1988). In each block, the same target patch was presented twice, once in the first half of the trials and once in the second half in randomized order. Response consistency is the proportion of trial pairs on which the observer gave the same response, regardless of the correctness (chance level=50%).

Before the main experiment, each observer completed a titration session using the same task at each location to determine the RMS contrast of the Gabor to maintain the overall performance at 70%. Gabor contrast was adjusted using a PEST method (Pentland, 1980) and averaged over two interleaving staircases each containing 50 trials.

After practice sessions, observers completed on average 4,317 trials (±262 trials) per location. All observers completed an equal number of trials per location, during 10-15 one-hour sessions. The Gabor contrast in the main experiment was adjusted according to the accuracy of the previous block: if the accuracy was higher/lower than 70%, the contrast was reduced/increased proportionally. Contrast sensitivity was calculated by taking the reciprocal of the contrast threshold of the last ten sessions in which titrated contrast thresholds were stable.

#### Eye-tracking

Eye position was monitored online using Eyelink1000. If the observer’s gaze deviated more than 1.5 dva away from the fixation center before the response, the trial was aborted immediately and repeated at the end of the block.

### Reverse correlation

Figure 2 illustrates the procedure of reverse correlation to estimate weights assigned by the system to different orientations and SFs (Ahumada, 2002; Li et al., 2016; Fernández et al., 2022). Reverse correlation enables experimenters to discretize the stimulus feature by varying the of sampling resolution in an efficient and flexible way. First, we quantified the energy profile (*E*_*θ,f*_) of the noise patch in the noise (*N*) by dot-multiplying the noise patch with a pool of Gabor-filters (*g*_*θ,f*_), defined by *θ* and *f*, in quadrature phase and have the same size of the target (**Fig 2A**, *Step 1)*. These Gabor filters approximate the linear filters in V1 that are tuned to a specific orientation and SF.

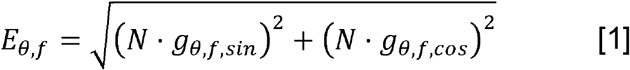

where θ stands for orientation, ranging from -90 to 90° in 29 steps relative to and centered at the Gabor orientation (0°), and *f* stands for SF, ranging from 1 to 4 cpd in 29 log steps, centered at the Gabor SF (2 cpd). No energy beyond 4 cpd was detected.

**Figure 2.**
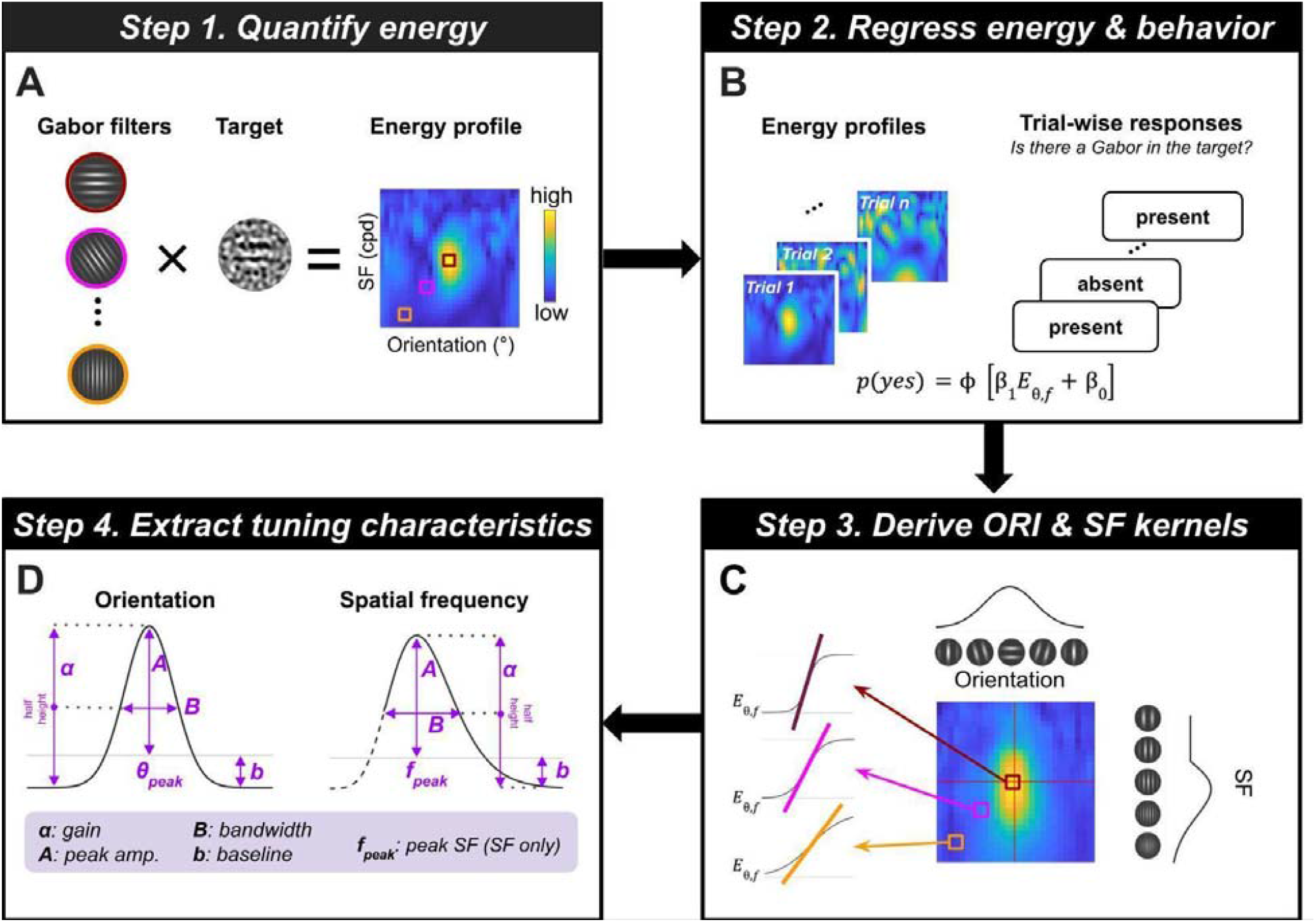
Reverse correlation. (**A**) Step 1. We computed energy profiles by dot-multiplying a pool of Gabor filters in quadrature phase with the target patch. (**B**) Step 2. We regressed the trial-wise energy fluctuations with behavioral responses and used the regression slope (β_1_) to index perceptual sensitivity. (**C**) Step 3. The derived β_1_ of all orientation-SF channels in Step 2 yielded a 2-D kernel mapping. Then, we took the mean (marginals) of the 2-D mapping to derive orientation and SF kernels. (**D)** Step 4. We fitted tuning functions to the kernels and characterized the tuning function by extracting tuning parameters (i.e., gain) and characteristics (i.e., peak amplitude, bandwidth and baseline for orientation and SF, and peak SF, see Table 1).

Second, we regressed the energy pixels of a certain orientation-SF component (concatenated across all trials) with the binary behavioral responses using a probit link function (**Fig 2B**, *Step 2*)

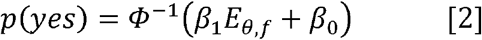

where *Φ*^−1^ (·)indicates the probit function (e., the inverse cumulative normal function), *β*_0_ is the intercept and the regression slope *β*_1_ is interpreted as the perceptual sensitivity to the corresponding orientation and SF. A more positive *β*_1_ indicates a higher sensitivity while *β*_1_ = 0 indicates no sensitivity. Before the regression, energy pixels at each orientation-SF channel across all trials were standardized (i.e., subtracting the mean across trials and divided by SD) for each Gabor contrast. We report results derived from Gabor-absent trials, which is a more conservative way to implement reverse-correlation analysis (e.g., Eckstein et al., 2002, 2004; Shimozaki et al., 2005).

**Table 1.** A list of tuning characteristics extracted from the orientation and SF tuning functions.

| For both orientation and SF |  |  |
| --- | --- | --- |
| Name | Computation | Interpretation |
| Gain ($\alpha$) | The scaling factor estimated by fitting a tuning function to the kernels | Overall sensitivity to the feature |
| Peak amplitude ($A$) | The height of the peak of the tuning function | Sensitivity to the task-relevant feature |
| Bandwidth ($B$) | Orientation: full width of function at half-height<br>SF: difference in log transform of SFs at half-height | Selectivity for the task-relevant feature |
| Baseline ($b$) | The height of the rightmost kernel | The baseline sensitivity |
| For SF only |  |  |
| Peak SF ($f_{peak}$) | The SF corresponding to the peak of the tuning function | The preferred SF of the system |

Third, we marginalized the kernel mapping across SF to obtain orientation sensitivity kernels and across orientation to obtain SF sensitivity kernels (**Fig 2C**, *Step 3*). Before this step, we assessed the separability between orientation and SF, calculated as the correlation (*r*^*2*^) between the raw kernel mapping and the reconstructed mapping (i.e., the outer product of the orientation and SF kernels), to ensure that marginalizing the 2D mapping would not lose feature information.

### Orientation and SF tuning functions

To capture the sensitivity to different orientations, we fitted a scaled Gaussian function to the orientation sensitivity kernels assuming that sensitivity kernels peaked at the Gabor orientation (i.e., 0°)

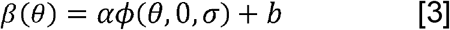

where *Φ* [·] stands for the normal probability function, *α* stands for the gain, *σ* stands for the Gaussian SD, and *b* stands for the baseline shift.

To capture the sensitivity to different SFs, we fitted a log parabola function to the SF sensitivity kernels (adapted from Watson & Ahumada, 2005).

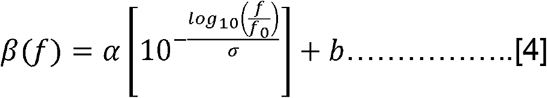

where *α, σ* and *b* indicates SF gain, standard deviation and baseline, respectively.*f*_0_ stands for the peak of the function. We used the MATLAB *fmincon* function to search for the parameters (*α, σ* and *b* for orientation and *α, σ, b*, and *f*_0_ for SF) using the Least-Square optimization method. Sensitivity kernels at the fovea and perifovea were fitted together. The parameter structure was varied by whether a parameter was fixed or free to vary between fovea and perifovea, yielding 8 candidate models for the orientation domain and 16 models for the SF domain. The best fitting candidate model was determined by model comparison with a 10-fold cross-validation method based on deviance averaged across 10 folds. The optimization procedure was repeated 20 times using the MATLAB *MultiStart* function with randomized initial parameters each time and the best fit was selected. Before testing the candidate models, we also compared two families for orientation (scaled Gaussian vs. difference of Gaussians (DoG)) and for SF (log parabola with vs. without truncation). In **Figure 7** we will present the model comparison results.

### Tuning characteristics

Finally, we extracted tuning characteristics (i.e., values characterizing the tuning function, illustrated in **Fig 2D**, *Step 4*) for comparison across locations. See **Table 1** for a complete list of tuning parameters and characteristics. We calculated the inter-parameter correlation which is the partial correlation between two parameters across observers while controlling for location. Compared to estimated parameters (e.g., gain and width), tuning characteristics (e.g., peak amplitude and bandwidth) have lower inter-parameter correlations, indicating higher independence among them and thus higher interpretability (**Figure 3**). Note that the gain and peak amplitude are two computationally different measures that reflect different tuning properties: Gain indicates the general amplitude amplification which depends on the baseline, whereas the peak amplitude indicates the highest sensitivity the system can achieve which is independent of the baseline.

**Figure 3.**
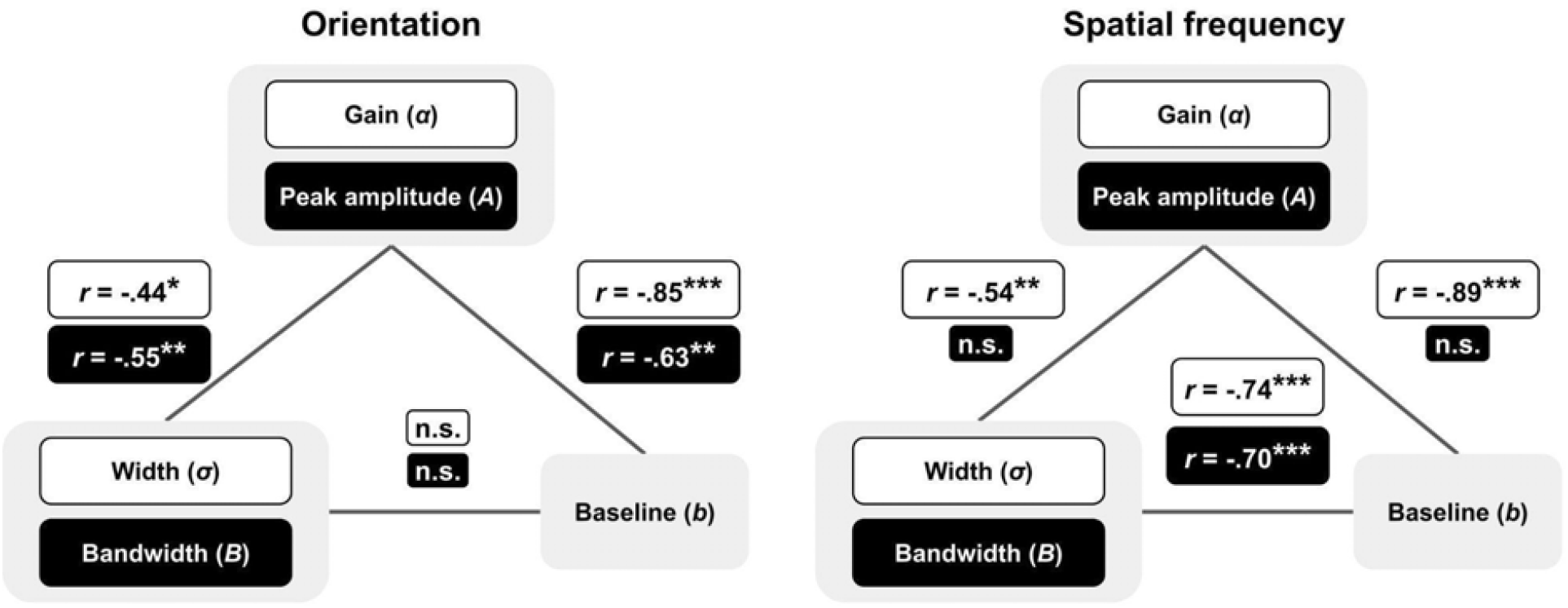
Correlations among estimated parameters and tuning characteristics derived from orientation (left panel) and spatial frequency (right panel) tuning functions across observers. The white boxes indicate estimated parameters and the black boxes indicate tuning characteristics. The symbols at the connecting lines indicate the strength of inter-parameter correlations. The pair being compared is indicated by the color of the box. (Note that the estimated baseline is equivalent to the extracted baseline). ***p < 0.001, **p < 0.01, *p < 0.05, n.s., p > 0.1.

Unlike many studies that only studied a single location or visual field (Hilz and Cavonius, 1974; Rovamo et al., 1978; Levi and Klein, 1986), we pooled four polar angles to represent the perifovea as they reflect the behavioral heterogeneity at perifovea (Carrasco et al., 2001; Barbot et al., 2021) so that we can generalize the revealed tuning properties.

### Statistical Analysis

After deciding the best fitting function via model comparison to characterize orientation and SF sensitivity kernels, we used a bootstrapping procedure to derive median and confidence interval (CI) of the tuning characteristics. In each iteration, we resampled the trials with replacement, regenerated sensitivity kernels using reverse correlation, and extracted tuning characteristics at the fovea and perifovea. After repeating this process for 1,000 times, we calculated and reported the median and 68% CI (which equals an SE, Wichmann and Hill, 2001) of all measurements and estimates for each observer. We then reported the mean and ±1 SEM across observers to represent the group level.

To compare performance (e.g., *d*-prime, criterion, contrast sensitivity and response consistency) and the computations (i.e., tuning characteristics and estimated internal noise) between fovea and perifovea, we conducted within-subjects ANOVAs and pair-wise two-tailed *t*-tests. When appropriate (e.g., when comparing tuning functions at each channel), we conducted Bonferroni’s correction to adjust for multiple comparisons.

### Noisy observer model

To assess the validity of the sensory representation and to estimate internal noise, we implemented a *noisy observer model* to predict trial-wise responses based on the *energy profile* of each trial and the *template* (i.e., the featural representation derived from Gabor-absent trials via reverse correlation). **Figure 4** illustrates the schematic of the model. **Table 2** lists all model parameters. Each iteration consisted of four steps. First, the data at each location were reshuffled and split into two sets (**Fig 4**, *step 1*). The first set was used to derive the template (*T*_*θ,f*_) using reverse correlation (as described above) and the second set was used to estimate free parameters by providing trial-wise energy profiles (*E*_*θ,f*_) and responses.

**Table 2.** Parameters in the noisy observer model.

| Model expressions |  | Model fitting |  |
| --- | --- | --- | --- |
| $T_{\theta,f}$ | Template estimated by applying reverse correlation to the *training* set | $n_{pc}$ | The total number of correct trials |
| $E_{i,\theta,f}$ | Energy profile of the $i$-th trial in the test set | $m_{pc}$ | The total number of trials |
| $V_i$ | The internal variable of the $i$-th trial | $Pc$ | The predicted proportion of correctness |
| $\mu_{S+}, \mu_{S-}$ | The mean of the internal variables for the Gabor-present and -absent trials, respectively | $n_{pa}$ | The total number of consistent trial pairs |
| $\sigma_{S+}^2, \sigma_{S-}^2$ | The variance of the internal variables for the Gabor-present and -absent trials, respectively | $m_{pa}$ | The total number of trial pairs |
| $V_{S+,noisy},$<br>$V_{S-,noisy}$ | The theoretical expression of noisy internal variables for the Gabor-present and -absent trials, respectively | $Pa$ | The predicted response consistency |

Table 2. Parameters in the noisy observer model.
| Free parameters |  |  |  |
| --- | --- | --- | --- |
| $N_{IE}$ | The ratio of the SD of induced noise to the external variability (i.e., $\sigma_{S+}^2$ and $\sigma_{S-}^2$) | $\sigma_{constant}^2$ | The variance of the constant noise |
| | | $k$ | The threshold |

**Figure 4.**
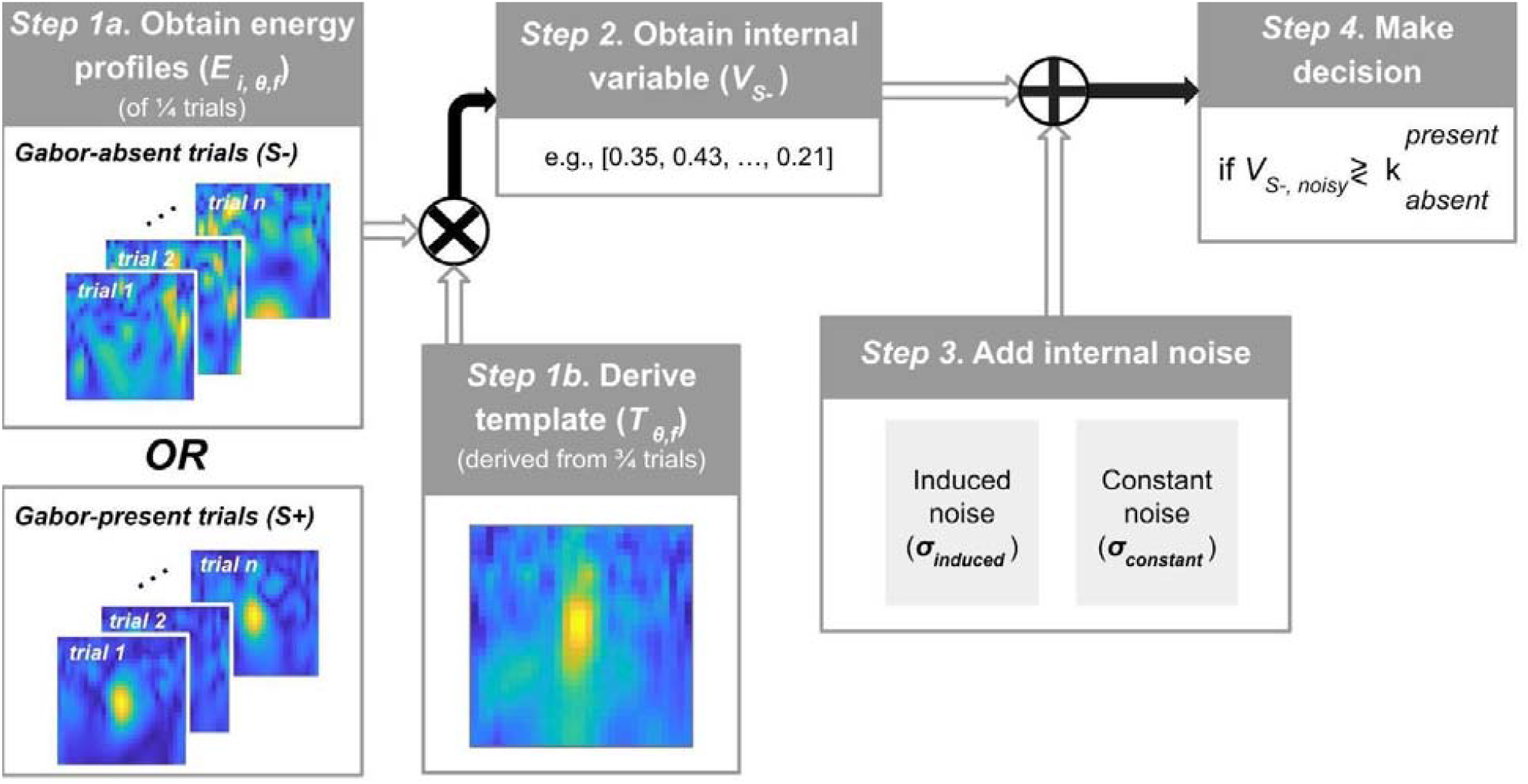
Schematic of the noisy observer model. **Step 1a**. A representation of orientations and SFs, which is assumed to be the template adopted by the observer to complete the detection task at one location, is derived from the training set (75% of trials per location). **Step 1b**. Energy profiles of the test set (25% trials) are calculated. **Step 2**. The internal variable for each trial is calculated by dot multiplying the template and the energy profile of each trial, separately for Gabor-present and -absent trials. **Step 3**. For each trial, a ‘noisy internal variable’ was created by adding two sources of internal noise to the internal variable. **Step 4**. A decision is made by comparing the noisy internal variable to a threshold.

Second, we calculated the *internal variable* (*V*_*i*_) for each trial in the test set by taking the sum of the dot product between the derived template and the energy profile of each trial (**Fig 4**, *step 2*)

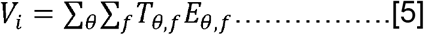

where (*V*_*i*_) stands for the internal variable of each trial. The internal variable forms the basis of trial-wise decision making, following the decision variable assumption of Pelli, (1985). Assuming that internal variables of all Gabor-present (*V*_*S* +_) and -absent trials (*V*_*S* −_)are normally distributed, means (μ_S+_ and μ_S −_) and variances (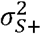 and 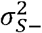) were extracted. Note that the integration of template and energy profile is a convolution in the spatial domain.

Third (**Fig 4**, *step 3*) we derived noisy internal variable (*V*_*i,noisy*_) by adding two types of internal noise to (*V*_*i*_): (1) *induced noise*, assumed to be drawn from a normal distribution whose width is proportional to the external variability (Burgess and Colborne, 1988; Pelli and Blakemore, 1990; Lu and Dosher, 2008; Diependaele et al., 2012) and (2) *constant noise*, whose variance is independent of external variability. Thus, noisy internal variables for Gabor-present and -absent trials, respectively, are characterized by a normal distribution

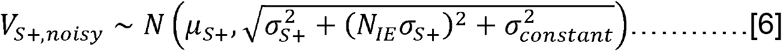

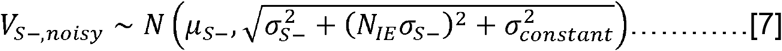

where *N*(·) stands for Normal distribution,*N*_*I E*_ is the internal-external noise ratio and 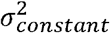 is the variance of the constant noise.

Finally (**Fig 4**, *step 4*), the response of this trial in the test set was determined by and comparing (*V*_*i,S+noisy*_) (or *V*_*i,S_noisy*_), to a threshold (*k*). The response is present if *V*_i_ ≥*k*. and absent if *V*_i_ < *k*

The model was fitted to each observer’s responses by minimizing the negative log-likelihood of correct responses consistent pairs (Lu and Dosher, 2008; Fernández et al., 2022),

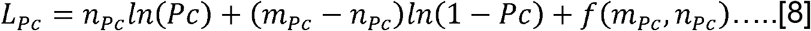

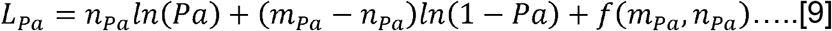

where *n*_*PC*_is the total number of correct trials, *m*_*PC*_ is the total number of trials, *PC* the predicted proportion of correctness, *n*_*Pa*_ is the total number of consistent trial pairs, *m*_*Pa*_ is the total number of trial pairs, and _*Pa*_ is the predicted response consistency.

*f* (*m*_*P*_, *n*_*P*_) is a function to calculate the factorial term in log space (Lu and Dosher, 2008)

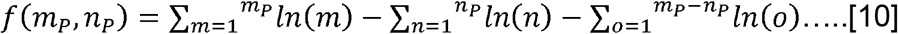

The expression of predicted response consistency is derived below. Take Gabor-present trials as an example, the response consistency is the proportion of Gabor-present trials (*S*^+^) when either the responses of two passes (*R*_A_ and *R*_B_) are both present (*R*_*A*_ = *R*_*B*_ = 1)or absent (*R*_*A*_ = *R*_*B*_ = 0)

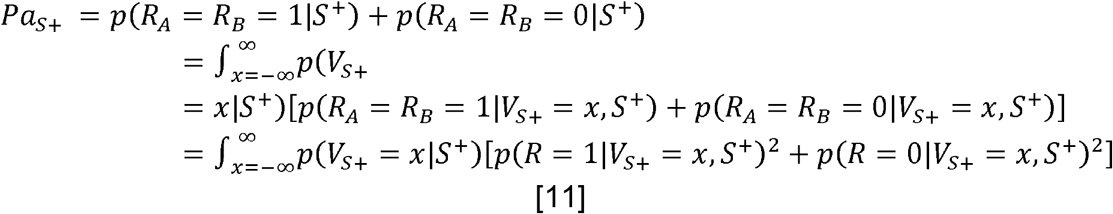

where

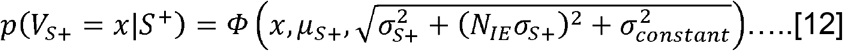

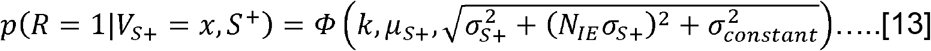

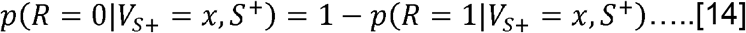

*x* is any possible values *V*_*S* +_ could take. Therefore, the predicted response consistency for Gabor-present and -absent trials are

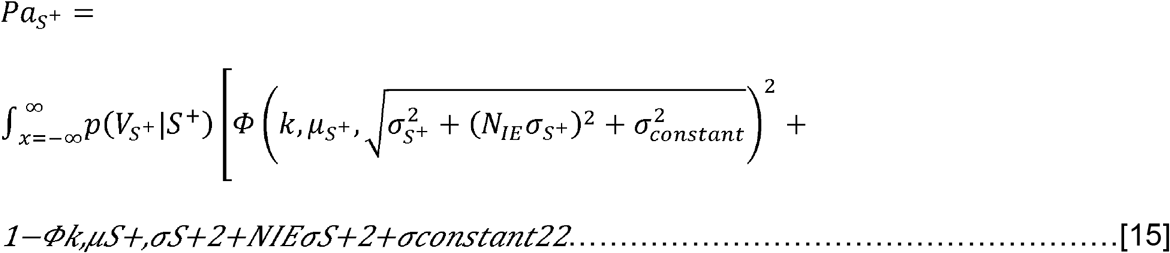

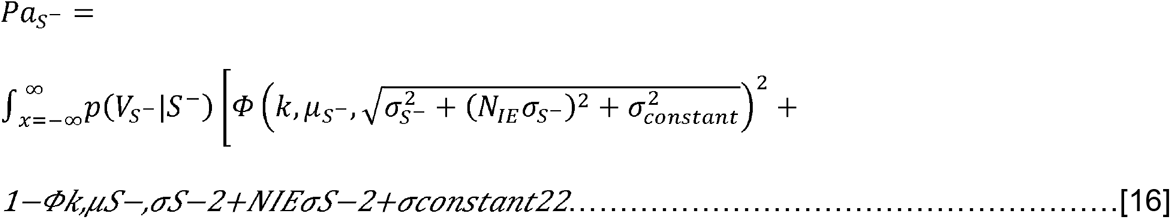

and the predicted response consistency of all trials is 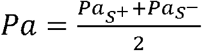

Finally, to evaluate the validity of the feature representation, we compared the predictive power of four model variations defined by whether the model contains any randomness (adapted from Beard and Ahumada, 1999):

1. *Core* model: neither the template nor the trial-wise energy profiles were random;
2. *Random template* model: pixels of the template were randomized on each trial;
3. *Random energy* model: pixels in each energy profile were randomized;
4. *Random template and energy* model: pixels in each energy profile and in the template were randomized.

The rationale is that a model without any randomness (i.e., to fully utilize the information of the template and energy profiles) should outperform other models. We calculated the Bayesian information criterion (BIC) score for each model variation at each location 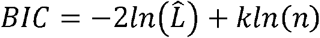 using where 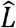 is the likelihood given the optimal parameter combinations of the candidate model, is the number of parameters in the candidate model and *n* is the number of data points. We reported differences in BIC scores Δ *BIC*_*i*_= *BIC*_*i*_ − *BIC*_*i,min*_ The best model is when Δ *BIC*_*i*_ ≡ Δ_*min*_≡ 0.

### Correlations between behavior and computations

To bridge behavior (i.e., contrast sensitivity) and computations (i.e., tuning characteristics and internal noise), we assessed the correlations between contrast sensitivity with tuning characteristics and with estimated internal noise across observers. To ensure that the correlations are not driven by possible location differences (e.g., higher orientation gain at fovea), we conducted a partial correlation analysis while controlling for the location effect.

Then, to further pinpoint the computation that underlies the eccentricity effect, we calculated the correlations between the eccentricity effect index (EEI):

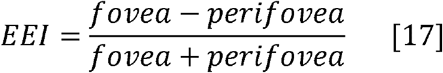

for contrast sensitivity, with tuning characteristics and with estimated internal noise. A positive index means that the assessed value at the fovea is higher than the perifovea and vice versa. A more positive or negative index indicates a larger difference between fovea and perifovea. To visualize the observer effect, we plot individual data with group averages across locations subtracted.

## Results

### *d*-prime and criterion are equated between the fovea and perifovea

**Figure 5A-** shows that there was no difference between fovea and perifovea in *d*-prime (t(11) = -1.02, p > 0.1) or criterion (t(11) = 1.17, p > 0.1), indicating matched performance across locations. This matched performance ensures that any change in the perceptual sensitivity is attributed to the trial-to-trial energy fluctuations rather than to task difficulty or the probability of detecting the target.

**Figure 5.**
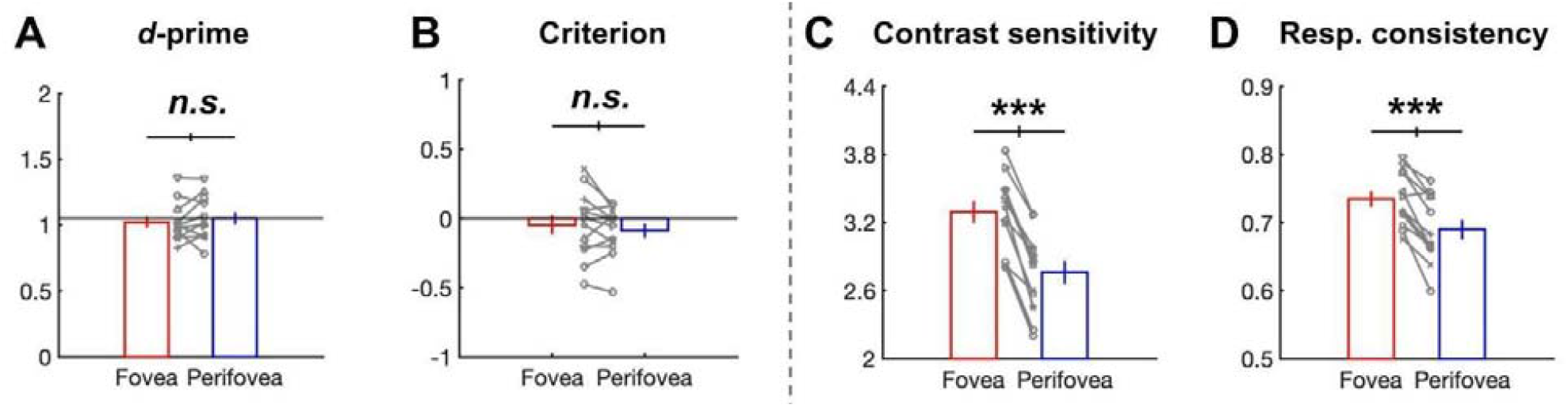
(A) d-prime, (B) criterion, (C) contrast sensitivity (reciprocal of contrast threshold) and (D) response consistency at the fovea and perifovea. Bars indicate group averages and error bars are ±1 SEM. Gray markers plotted between two bars represent individual data (the median of 1K bootstrapping) and each observer is indicated by a unique marker type. Error bars representing ±1 SEM. The error bars intersecting the black horizontal line indicate ±1 SEM of the difference. ***p < 0.001, n.s., p > 0.1.

### Higher contrast sensitivity and higher response consistency at the fovea

**Figure 5C and 5D** show the contrast sensitivity and response consistency measured using a double-pass method (Burgess and Colborne, 1988). Contrast sensitivity was higher at fovea than perifovea (t(11) = 10.07, p < 0.001), confirming the established eccentricity effect, which was present for all observers. There was also a higher response consistency at the fovea than perifovea (t(11) = 6.06, p < 0.001; true for all but one observer for whom there was no difference), suggesting lower internal noise in the system at the fovea (Burgess and Colborne, 1988). This response consistency measurement is the key to directly estimating the level of internal noise via the implementation of a noisy observer model.

### Compare 2D kernel mapping

The sensitivity kernel mapping at the fovea and perifovea was derived using reverse correlation from Gabor-absent trials only (**Figure 6A-B**). Each pixel indicates the perceptual sensitivity, i.e., how influential those two channels are in the perception of the stimuli at a certain orientation and SF channel. **Fig 6C** shows that the most pronounced positive differences between the fovea and perifovea were found near the Gabor features (represented by the red horizontal and vertical lines), suggesting that the fovea is more sensitive to the task-relevant features. **Fig 6D** shows the ideal template which is the energy profile of the signal Gabor. We confirmed that orientation and SF dimensions are separable for most observers (**Fig 6E**). The higher the correlation, the more separable the two dimensions are and the less information is expected to be lost by examining the two dimensions separately. High separability justifies marginalization along each dimension to derive orientation and SF tuning functions (**Figure 7**).

**Figure 6.**
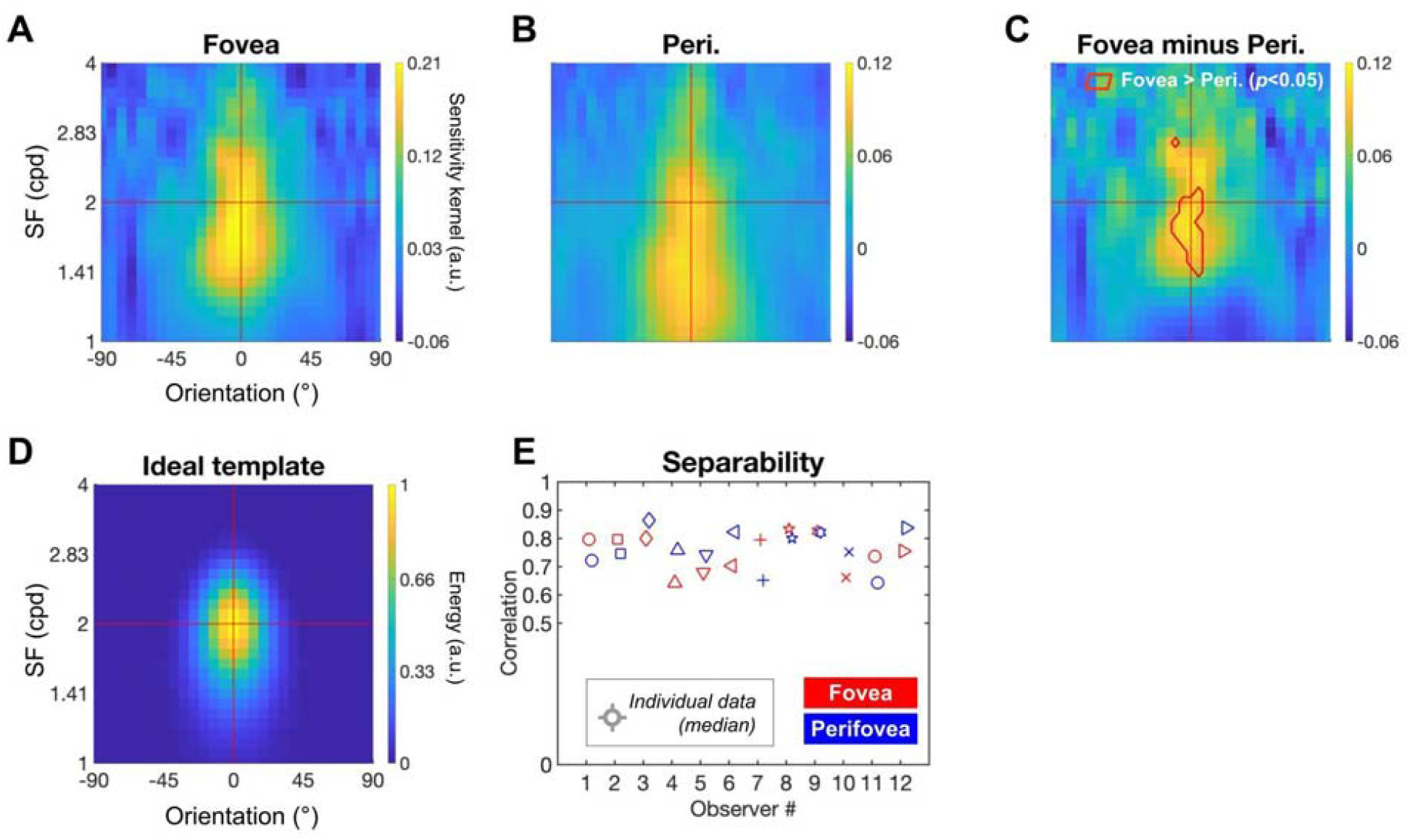
2-D sensitivity kernel mapping and separability derived from Gabor-absent trials. Pixels in A-C represent group-averaged sensitivity kernels derived from data at the fovea (**A**) and perifovea (**B**), indicating the regression slope representing the correlation between the feature energy and the behavioral responses. The mapping’s horizontal and vertical red lines depict the Gabor SF and orientation, respectively. (**C**) Difference between fovea and perifovea. The red border indicates the orientation-SF channel at which the fovea is higher than the perifovea (p < 0.05, Bonferroni corrected one-way t-test). (**D**) The ideal template which is the energy profile of the signal derived from step 1 of **Fig 2**. (**E**) Separability (i.e., the correlation between reconstructed sensitivity kernels and original kernels). A correlation close to 1 implies high separability.

**Figure 7.**
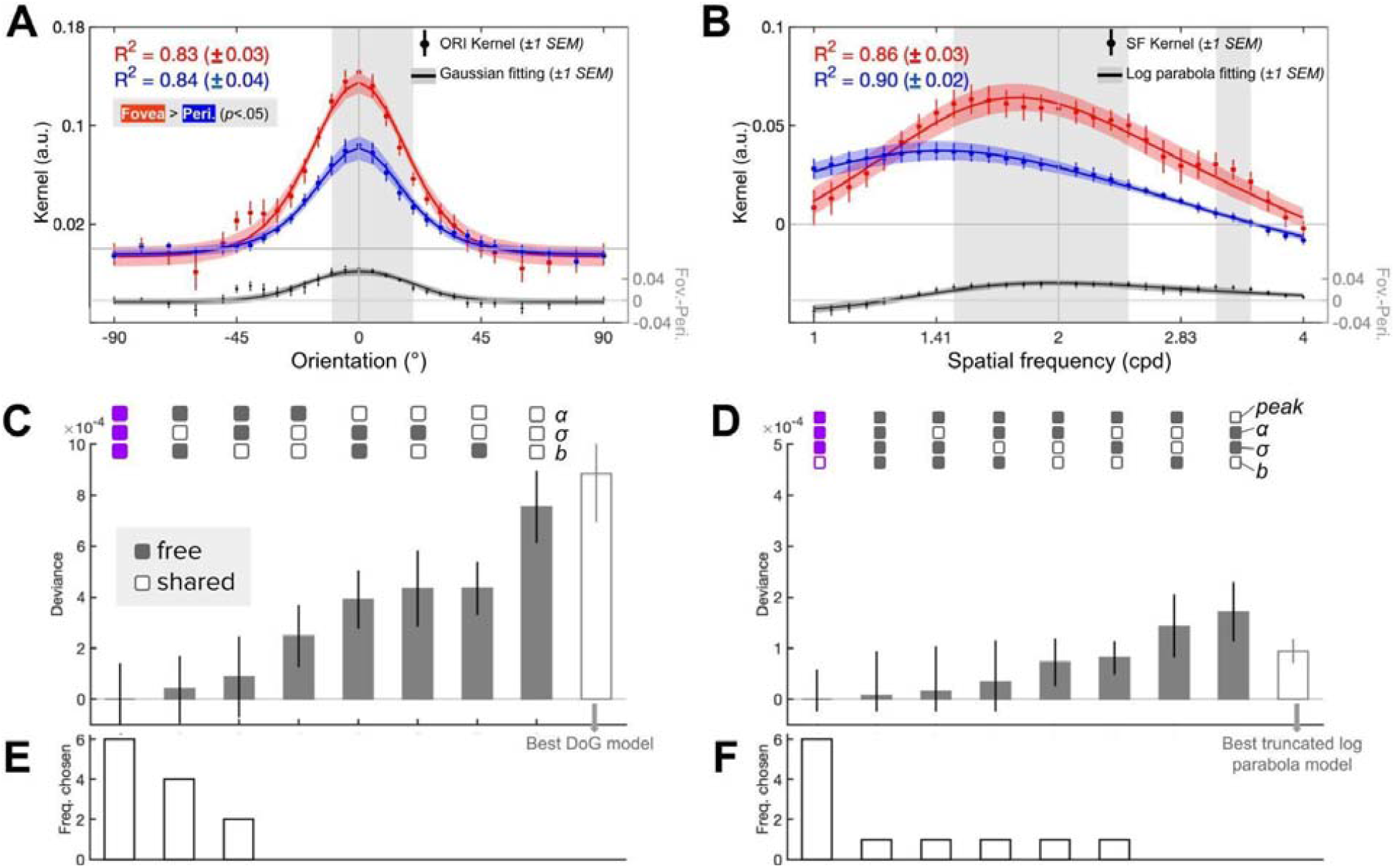
Marginalized kernels and the sensory tuning function for orientation (**A**) and SF (**B**) with model comparison (**C-D**). (**A and B**) Red (fovea) and blue (perifovea) dots are group-averaged marginalized kernels and error bars represent ±1 SEM. Curves with corresponding colors are group-averaged sensory tuning functions and bands represent ±1 SEM. Asterisks indicate channels at which the fovea is higher than the perifovea (Bonferroni-corrected post-hoc one-tailed t-test, p < 0.05). Black dots below asterisks are the averaged difference between fovea and perifovea (indicated by the y-axis on the right) and bands in black are ±1 SEM. (**C and D**) Comparison among candidate models of the winning family (gray bars, scaled Gaussian for orientation and log parabola for SF), and with the best-fitting model of other families (white bar with gray outline, DoG for orientation and truncated log parabola for SF). Deviance is the difference between data and prediction using 10-fold cross-validation averaged across 10 folds. Empty squares: free to vary; filled-in squares: shared. (**E and F**) The frequency of each model being the best-fitting model when fitted to individual data.

### Comparing orientation and SF tuning functions

Figure 7. shows the marginalized kernels and the fitting functions for orientation (**Fig 7A**, red dots) and SF (**Fig 7B**, blue dots). First, we conducted a within-subjects two-way ANOVA (fovea vs. perifovea x 29 channels) on the marginalized orientation and SF kernels separately. There was a main effect of the channel for both orientation (F(28,308) = 54.00, p < 0.001) and SF (F(28,308) = 16.93, p < 0.001). Location had a main effect for both orientation (F(1,11) = 68.37, p < 0.001) and SF (F(1,11) = 133.32, p < 0.001); kernels were higher at fovea than perifovea across channels. Bonferroni-corrected post-hoc one-tailed *t*-tests (indicated by grey bands in **Fig 7A-B**, p < 0.05) revealed foveal advantage over perifovea near the task-relevant orientation and SFs and at higher SFs.

Then, to identify the tuning mechanisms, we captured the shape and magnitude of the marginalized kernels by fitting a scaled Gaussian function to the orientation kernels (**Fig 7A**, red curves) and a log parabola function to the SF kernels (**Fig 7B**, blue curves). Model comparisons revealed that orientation kernels were best fitted by the candidate model with all parameters free to vary (**Fig 7C**) and SF kernels by the candidate model with only baseline (*b*) fixed (**Fig 7D)**. The two best candidate models fit the data well (R^2^ = 83.5% for orientation and R^2^ = 88% for SF on average), allowing us to extract tuning characteristics from the fitting function as a proxy of tuning properties (**Fig 2D** and **Table 1**).

Comparing tuning characteristics between the fovea and perifovea showed that for orientation (**Fig 8, top row**), there was a higher gain (t(11) = 5.62, p < 0.001) and higher peak amplitude (t(11) = 7.79, p < 0.001) at fovea than perifovea, but no difference in bandwidth or baseline (p > 0.1). The differences in orientation gain and peak were highly consistent across individual observers. The results indicate that observers were more sensitive to orientation in general, especially to the task-relevant and neighboring orientations at fovea than perifovea.

**Figure 8.**
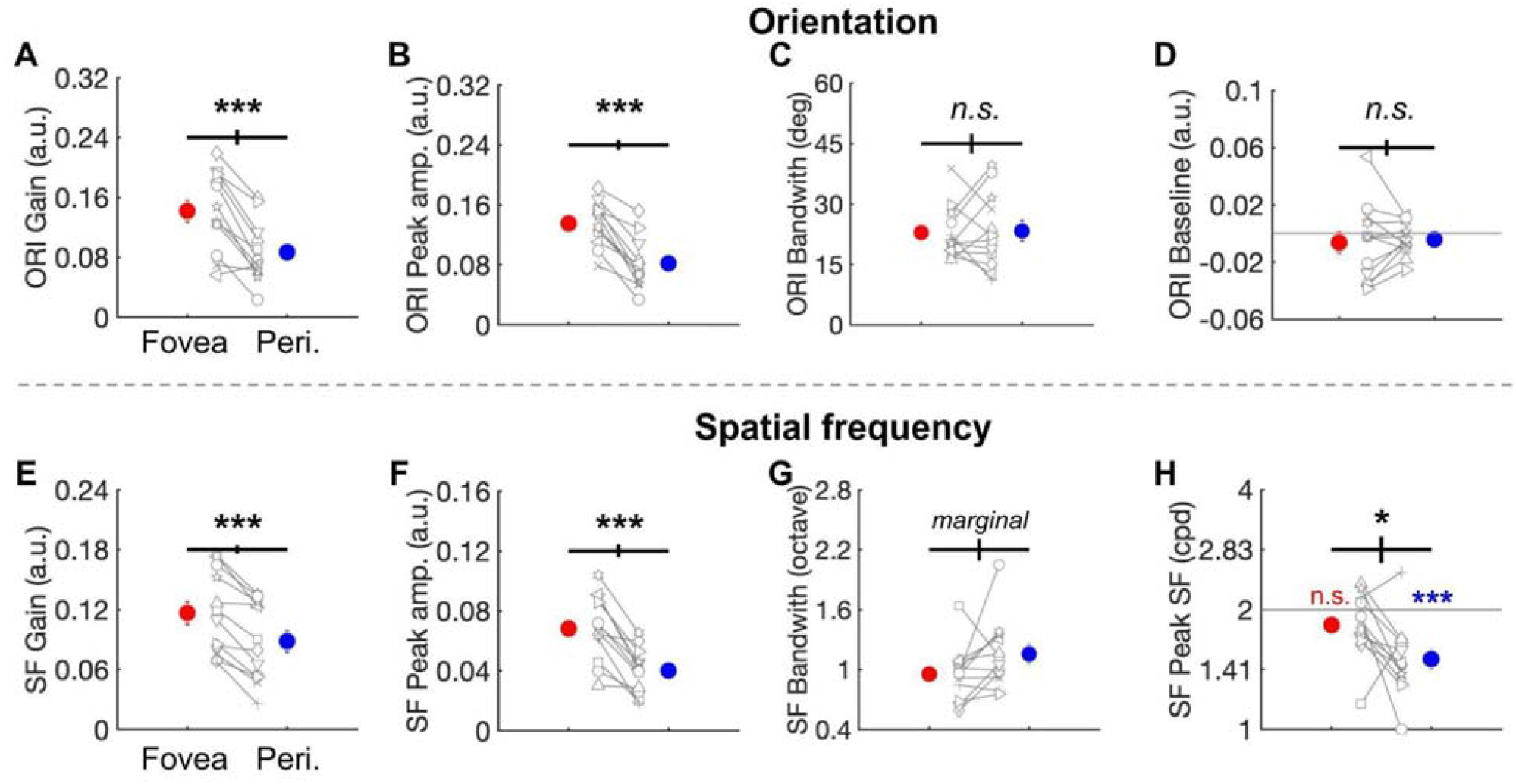
Comparison of orientation (**A-D**) and SF (**E-H**) tuning characteristics between fovea (red dots) and perifovea (blue dots). Dots represent group averages with ±1 SEM. Gray dots indicate individual data. ***p < 0.001, marginal, p > 0.05 and p < 0.1, n.s., p > 0.1.

For SF, there was a higher gain (**Fig 8E**, t(11) = 6.55, p < 0.001) and a higher peak amplitude (**Fig 8F**, t(11) = 6.39, p < 0.001) at fovea than perifovea, indicating that observers were more sensitive to SFs in general, especially to the task-relevant SFs. There was a marginal difference in SF bandwidth in octave (**Fig 8G**, t(11) = -1.83, p = 0.094), 9 out of 12 observers showed narrower SF bandwidth at the fovea. Given the presence of one outlier at each fovea and perifovea, we calculated both mean (fovea: 0.95, perifovea: 1.15) and median (fovea: 0.95, perifovea: 1.09), and the statistical results do not change. Finally, tuning functions peaked at a higher SF at fovea than perifovea (**Fig 8H**, t(11) = 2.57, p = 0.026); this was the case for 10 out of 12 observers. Moreover, the peak of SF tuning function at the fovea did not differ from the signal SF (2 cpd, indicated by the gray horizontal line in **Fig 8H**, p > 0.1) whereas the peak at the perifovea was lower than the signal SF (p < 0.001).

### Estimate internal noise via computational modeling

We could infer that fovea has lower internal noise because we observed a higher response consistency at the fovea than perifovea (**Fig 5**). To obtain the numerical estimation of internal noise, we implemented a noisy observer model to trial-wise responses by modeling the internal response and induced and constant internal noise (**Fig 4**). First, the predicted accuracy (**Fig 9A**) and response consistency (**Fig 9B**) match the data well; most dots lie near the diagonal (p > 0.1), indicating that the model has good predictive power. Second, response consistency negatively correlated with induced noise (**Fig 9C**, partial r = -0.80, p < 0.001) and constant internal noise (**Fig 9D**, partial r = -0.70, p < 0.001) as expected (Burgess and Colborne, 1988). Third, most observers showed lower induced noise (10 out of 12, t(11) = -2.50, p = 0.029, **Fig 9E**) and constant noise (9 out of 12, p > 0.1, **Fig 9F**) at the fovea than perifovea. In sum, the noisy observer model successfully predicted trial-wise responses and found constant internal noise lower at the fovea than perifovea.

**Figure 9.**
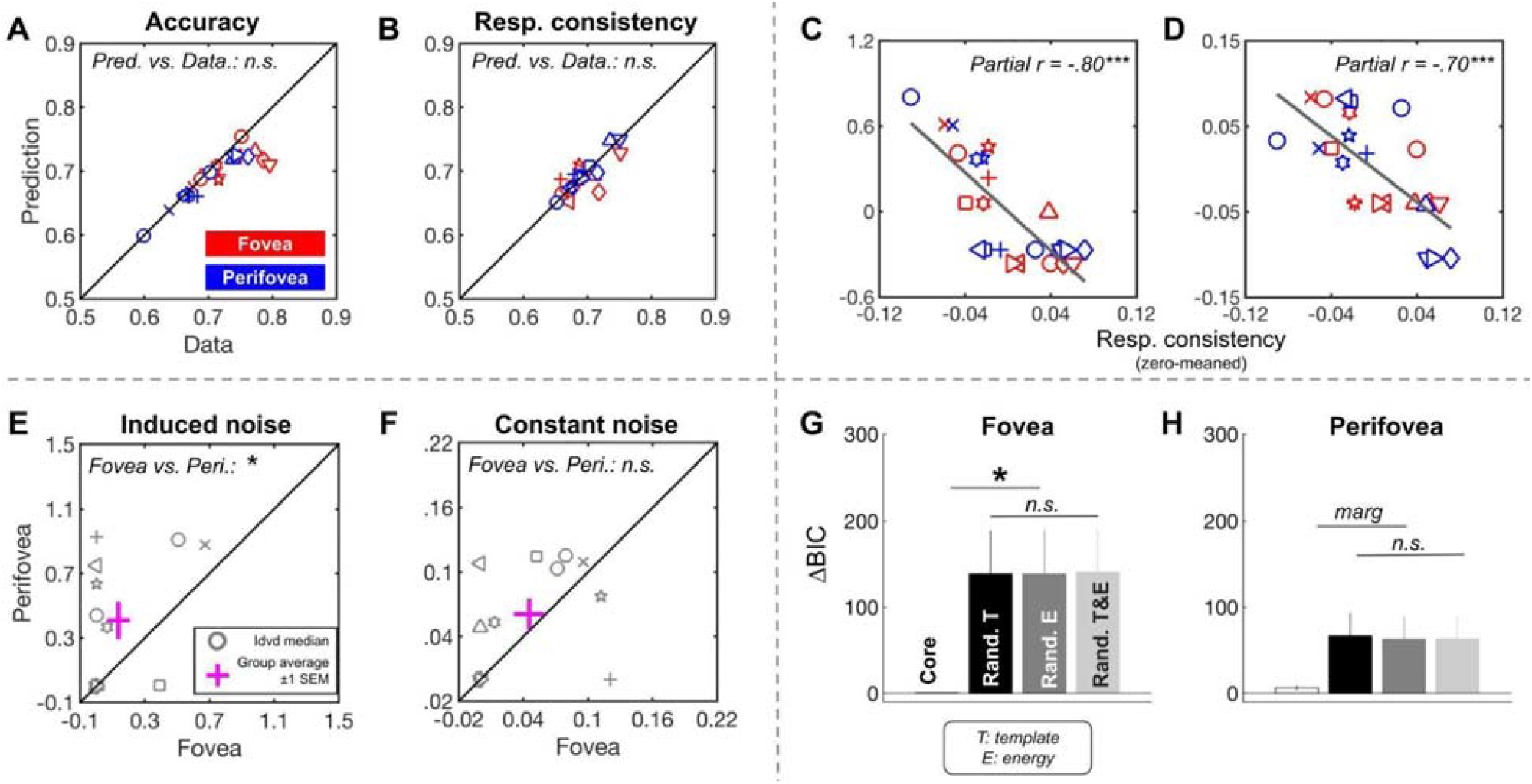
Evaluate noisy observer model. (**A-B**) Comparison of the predicted accuracy and response consistency (**B**) with data. (**C-D**) Correlation between the response consistency with the induced noise (**C**) and constant noise (**D**). (**E-F**) Comparison of the induced noise (**E**) and constant noise (**F**) between fovea and perifovea. Gray markers indicate individual data. (**G and H**) compare the predictive power (ΔBIC) of four model variations: the Core mode (white bar), the Random template model (black bar), the Random energy model (dark gray bar) and the Random template and energy model (light gray bar) for fovea (**G**) and perifovea (**H**). Bars represent group averages and error bars are ±1 SEM. ***p < 0.001.*p < 0.05, n.s., p > 0.1.

To assess the validity of the template (i.e., the representation for orientation and SF derived from partial data using reverse correlation), we compared the predictive power of four model variations (**Figs 9G-H**). The Core model outperformed all three random models (p = 0.019/0.019/0.016 (fovea) and p = 0.056/0.062/0.068 (perifovea), respectively). The three random models did not differ in their predictive power (p > 0.1). The results indicate that featural representation is crucial to predicting trial-wise responses.

### Linking tuning characteristics with contrast sensitivity

So far we have shown that feature representation differs between fovea and perifovea in sensitivity to and selectivity for orientation and SF. To bridge behavior and computations, we assessed the partial correlation between contrast sensitivity with tuning characteristics and with internal noise across observers (**Figure 10**). Contrast sensitivity correlates (1) positively with orientation gain (**Fig 10A;** partial r = 0.68, p < 0.001) and with orientation peak amplitude (**Fig 10B;** partial r = 0.75, p < 0.001), indicating that higher contrast sensitivity is associated with higher sensitivity to task-relevant orientations, (2) negatively with orientation bandwidth (**Fig 10C**; partial r = -0.41, p = 0.049) and with SF bandwidth (**Fig 10F**; partial r = -0.50, p = 0.016), indicating that higher contrast sensitivity is associated with higher selectivity for task-relevant features, (3) negatively with orientation baseline (**Fig 10D**, partial r = -0.42, p = 0.047), indicating that high contrast sensitivity is associated with the inhibition on orientations that are distant to the signal orientation, which is task-irrelevant (reflected by low or negative values in the inset of **Fig 10D**).

**Figure 10.**
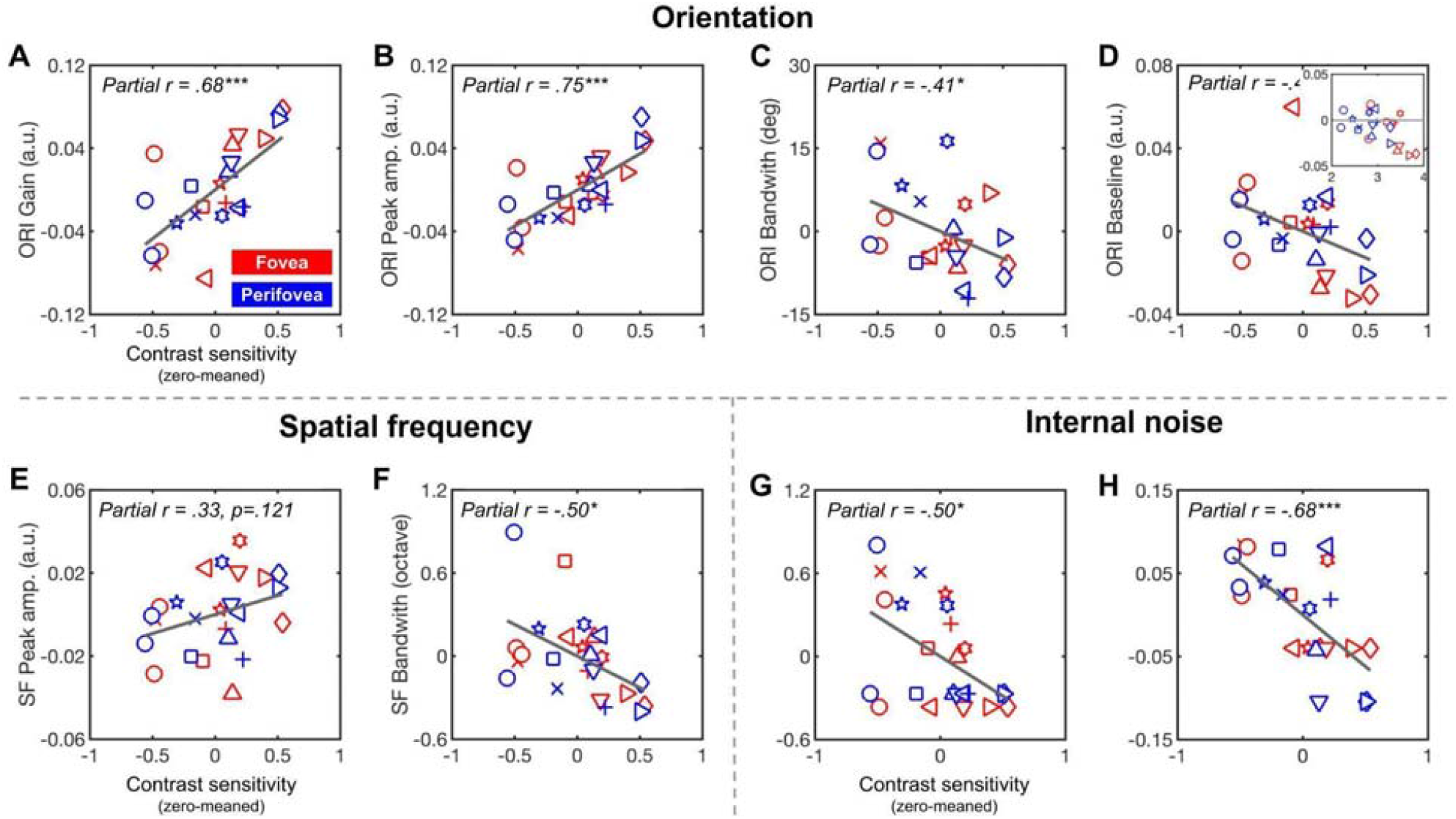
Correlation between contrast sensitivity and orientation tuning characteristics (**A-D**), SF tuning bandwidth (**E and F**), and estimated internal noise (**G and H**). Values in the figure were zero-meaned to reveal the observer effect. Markers are the median of individual data at the fovea (red) and perifovea (blue). Each individual data is represented by a unique marker. The black diagonal line is the linear regression line.

Then, we assessed the correlation between contrast sensitivity and internal noise. There was a negative correlation between contrast sensitivity and both induced noise (**Fig 10G**; partial r = -0.50, p = 0.014) and constant noise (**Fig 10H**; partial r = -0.68, p < 0.001), indicating that better detection was associated with lower internal noise.

Figure 11. shows the association between the eccentricity-effect index EEI for contrast sensitivity and computations. We found that an increase in the EEI of contrast sensitivity is associated with a stronger foveal advantage in the orientation peak amplitude (**Fig 11A**, r = 0.76, p = 0.004) and with narrower orientation bandwidth at the fovea (**Fig 11B**, r = -0.79, p = 0.002). No other correlations between EEI for contrast sensitivity and tuning characteristics were significant. These results indicate that the sensitivity to and selectivity for orientation play important roles in the underlying perceptual computations that give rise to the eccentricity effect. However, the EEI of internal noise does not correlate with the behavioral eccentricity effect (**Fig 11C and D**, p > 0.1), suggesting that differences in internal noise between fovea and perifovea do not predict the extent of the eccentricity effect in contrast sensitivity.

**Figure 11.**
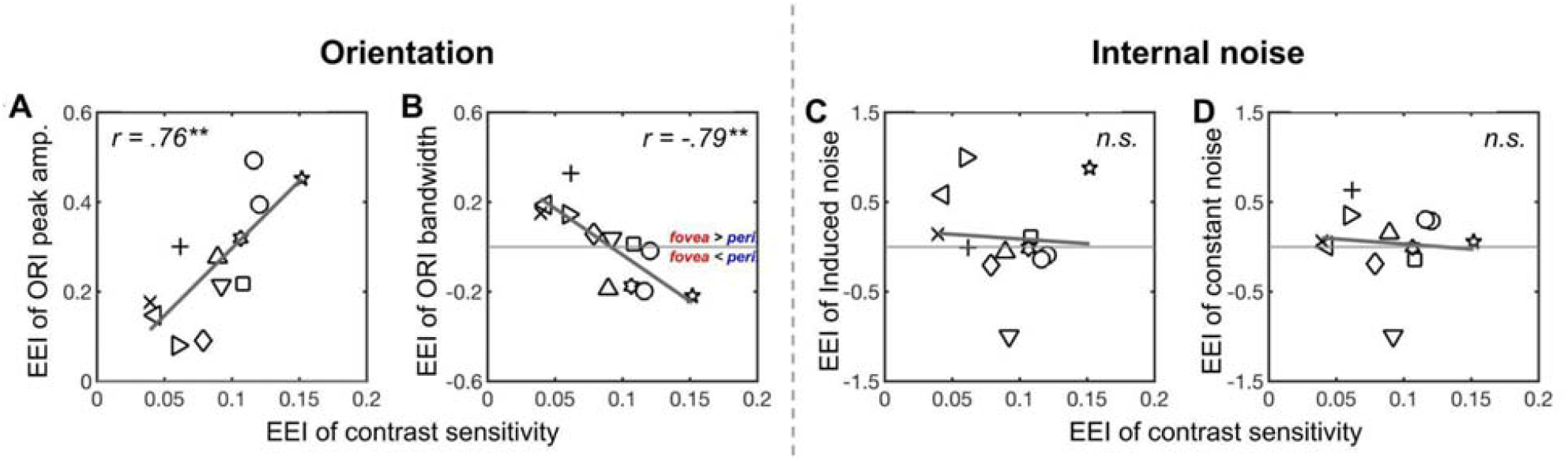
Correlation between the eccentricity effect index (EEI) of contrast sensitivity and (**A**) orientation gain (peak amplitude yielded the same result), (**B**) orientation bandwidth, (**C**) induced noise, and (**D**) constant noise. A positive EEI indicates fovea being higher than perifovea and vice versa. Greater positive/negative EEI indicates a larger fovea-perifovea difference. Markers are the median of EEI for each individual observer. The diagonal black line is the linear regression line. **p < 0.01, *p < 0.05, n.s., p > 0.1

## Discussion

We investigated whether performance differences between the fovea and perifovea, which result in an eccentricity effect (Rijsdijk et al., 1980; Carrasco et al., 1995, 1998; Carrasco and Frieder, 1997; Pointer and Hess, 1989; Baldwin et al., 2012), can be accounted for by distinct system-level computations –featural representation and internal noise. Using reverse correlation and the double-pass method, we established a link between performance and system-level computations and found that the behavioral advantage at the fovea stems from a better representation of task-relevant orientations and SFs, as well as lower internal noise.

### Featural representations differ between fovea and perifovea

#### Orientation sensitivity

We observed a higher orientation gain and peak amplitude at the fovea than perifovea indicating stronger sensitivity to task-relevant orientation at the fovea. Possible neural correlates are the higher density of retinal ganglion cells (Wässle et al., 1989) and higher neuron count corresponding to the fovea with neuron density approximately uniform across V1 (Hubel and Wiesel, 1977; Rockel et al., 1980).

#### Orientation selectivity

We found no difference in orientation bandwidth (fovea: 19.49°±1.65°, perifovea: 19.95°±2.35°), indicating a similar level of orientation selectivity. The comparable orientation bandwidth may seem surprising because the receptive field (RF) size of V1 neurons increases with eccentricity (Hubel and Wiesel, 1962) and V1 neurons with smaller RF have higher orientation selectivity (Watkins and Berkley, 1974). However, whereas complex V1 neurons’ orientation selectivity decreases with RF size, simple V1 neurons’ orientation selectivity is either less (Watkins and Berkley, 1974) or not (Schiller et al., 1976b) correlated with RF size, and featural representation estimated by reverse correlation approximates activity of simple V1 neurons (Neri and Levi, 2006). Indeed, a similar orientation bandwidth has been reported at 10° eccentricity in other reverse correlation studies (Li et al., 2016; Fernández et al., 2019). These findings are consistent with similar orientation bandwidth on average of simple V1 neurons at fovea and parafovea in monkeys (De Valois et al., 1982b; Xu et al., 2007). Note, however, that although orientation selectivity did not differ in our detection task, it may differ in a discrimination task.

#### SF sensitivity

We found (1) higher gain and peak amplitude at the fovea than perifovea, indicating higher sensitivity to the signal SF at the fovea, and (2) that SF tuning function peaked near the signal SF (2 cpd) at the fovea (1.86±0.09 cpd) but at a lower SF at the perifovea (1.53±0.10 cpd), indicating a better task-relevant SF representation at the fovea. These findings are consistent with a classification imaging study in which observers assigned more weight to lower SFs at the parafovea (1.25° and 2.5° eccentricity) than at the fovea (Levi and Klein, 2002). Similarly, reverse correlation yielded a lower SF peak than the signal SF at 10° eccentricity (Li et al., 2016; Fernández et al., 2019). Together, these studies indicate that the eccentricity effect in featural representation varies from fovea to parafovea and perifovea.

The finding that lower SFs are represented as more influential with increasing eccentricity is consistent with: (1) V1 simple cells preference for lower SF as eccentricity increases in macaques (De Valois et al., 1982b; Schiller et al., 1976a) and cats (Movshon et al., 1978a), (2) more cortex devoted to higher SFs in the fovea than parafovea (Xu et al., 2007), and (3) human contrast sensitivity function peaking at a higher SF at the fovea than perifovea (Robson and Graham, 1981; Wright and Johnston, 1983; Jigo et al., 2023). Preference for higher SF at the fovea may reflect the smaller RF of V1 simple cells at the fovea (Hubel and Wiesel, 1962), which yields a preference for higher SF (Movshon et al., 1978a, 1978b). Similarly, according to fMRI human studies, V1 voxels’ sensitivity to the signal SF decreases with eccentricity (Henriksson 549 et al., 2008; Aghajari et al., 2020).

Importantly, the mismatch between the SF tuning functions at the perifovea and the signal SF indicates that the task demand for this detection task cannot override the preference of lower SFs at the perifovea. This finding questions the assumptions of the linear amplifier (Pelli, 1981; Lu and Dosher, 2008) and perceptual template (Lu and Dosher, 1998, 2008, 2023) models, which match the template to the signal without considering visual field location. In the future, these and other (e.g., Lago et al., 2021) models should include the differential tuning features across eccentricity.

#### SF selectivity

We found similar SF bandwidth at the fovea (median: 0.95±0.08 octave) and perifovea (median: 1.09±0.10 octave), indicating comparable selectivity for task-relevant SFs at these locations. This estimation agrees with the typical 1-octave estimation of neurons (Blakemore and Campbell, 1969; Jakobsson and Lennerstrand, 1985) in human V1. There are mixed findings regarding SF selectivity across eccentricity. Similar SF bandwidth across eccentricity has been reported for V1 neurons in monkeys (De Valois et al., 1982a; Foster et al., 1985) and V1 voxels in humans (Broderick et al., 2022). But another fMRI study reported that the SF bandwidth of V1 voxels increases with eccentricity (Aghajari et al., 2020). This discrepancy might relate to the task design; whereas Broderick et al (2022) used stimuli that span a broad range of orientations and SFs, Aghajari et al. (2020) used stimuli defined by narrow SFs.

### Internal noise differs between fovea and perifovea

Our findings indicate that lower internal noise is another important factor underlying the performance advantage at the fovea. First, we inferred the level of internal noise from the response consistency using the double-pass method measured at the fovea (73%±1%) and perifovea (69%±1%). These values are similar to previous reports at the fovea (Burgess and Colborne, 1988; Murray et al., 2002; Ratcliff et al., 2018; Vilidaite and Baker, 2017).

Then, we directly estimated the level of internal noise using a noisy observer model for induced (fovea: 0.14±0.07; perifovea: 0.41±0.12) and constant (fovea: 0.05±0.01; perifovea: 0.06±0.01) noise. Our estimation of lower induced noise than previous reports at the fovea (0.65-1.3; Burgess and Colborne, 1988; Levi and Klein, 2003; Gold et al., 2004) is likely due to: (1) their estimate being based on measuring both types of internal noise with a common parameter; (2) internal noise estimates using static stimuli varying in basic features or yes-no detection tasks, like the current study, tend to fall on the lower end of the distribution (0.2-0.5; Neri, 2010).

We found internal noise lower at the fovea than perifovea for the basic features of orientation and SF. For a fixed-size stimulus, more neurons at the fovea may average out the internal noise (Kara et al, 2000). This finding is consistent with varying internal noise with eccentricity for motion (Mareschal et al., 2008a) and depth (Falkenberg and Bex, 2007; Wardle et al., 2012). However, there is similar internal noise across eccentricities in a face identification task (Mäkelä et al., 2001). Internal noise levels across eccentricity may differ for tasks that rely on distinct cortical areas.

Spatial uncertainty is higher in the periphery than fovea for localization precision tasks when spatial uncertainty is induced (Hess and Hayes, 1994; Michel & Geisler, 2011). This difference in spatial uncertainty is unlikely to contribute to the observed eccentricity effect or location differences in the tuning properties, because our design minimized spatial uncertainty by (1) adopting a detection task that does not require high spatial precision; (2) blocking target location and informing observers of the 100% valid target location; (3) having one white and three black placeholders indicating the target and non-target locations, respectively, throughout the trials.

### Differential computations mediate behavioral differences between fovea and perifovea

This is the first study to link performance and two system-level computations –featural representation and internal noise– in humans. First, a correlation analysis revealed that higher contrast sensitivity at fovea than perifovea can be attributed to (1) a better representation of task-relevant features and (2) less internal noise in the system. Correlations have been established between neural structure with performance (Duncan and Boynton, 2003; Song et al., 2015; Himmelberg et al., 2022) and with computations (Neri and Levi, 2006). Thus, our findings on the correlation between performance and computations can inform the interaction among behavior, computation and neural structure. Second, we tested the hypothesis that the eccentricity effect is reflected in the differential computations between fovea and perifovea. Our analyses revealed that the greater the advantage in sensitivity to and selectivity for task-relevant orientations at the fovea than perifovea, the stronger the observed eccentricity effect. This result allows us to pinpoint orientation sensitivity and selectivity as dominant computations underlying the behavioral eccentricity effect in this detection task.

To conclude, we found (1) observers are more sensitive to orientations and SFs at fovea than perifovea, similarly selective for orientations and SFs at fovea and perifovea, and shift their peak sensitivity to lower-than-signal SFs at the perifovea, (2) internal noise is lower at fovea than perifovea, (3) a link between individual differences in contrast sensitivity with both feature tuning characteristics and internal noise. Together, these findings provide compelling evidence that the foveal advantage in visual tasks can be attributed to two system-level computations: a better featural representation and lower internal noise than at perifoveal locations.

## Acknowledgements

This research was supported by National Institutes of Health Grants R01-EY027401(M.C.).

## References

Abbey, C. K., Eckstein, M. P. (2002). Optimal shifted estimates of human-observer templates in two-alternative forced-choice experiments. IEEE Transactions on Medical Imaging, 21(5), 429–440.

Abbey, C. K., & Eckstein, M. P. (2009). Frequency tuning of perceptual templates changes with noise magnitude. Journal of the Optical Society of America. A, Optics, Image Science, and Vision, 26(11), B72–B83.

Aghajari, S., Vinke, L. N., Ling, S. (2020). Population spatial frequency tuning in human early visual cortex. Journal of Neurophysiology, 123(2), 773–785.

Ahumada, A. J. (2002). Classification image weights and internal noise level estimation. In Journal of Vision (Vol. 2, Issue 1, p. 8). 10.1167/2.1.8

Anton-Erxleben, K., Carrasco, M. (2013). Attentional enhancement of spatial resolution: linking behavioural and neurophysiological evidence. Nature Reviews. Neuroscience, 14(3), 188–200.

Baldwin, A. S., Meese, T. S., Baker, D. H. (2012). The attenuation surface for contrast sensitivity has the form of a witch’s hat within the central visual field. Journal of Vision, 12(11). 10.1167/12.11.23

Barbot, A., Xue, S., Carrasco, M. (2021). Asymmetries in visual acuity around the visual field. Journal of Vision, 21(1), 2.

Benson, N. C., Kupers, E. R., Barbot, A., Carrasco, M., Winawer, J. (2021). Cortical magnification in human visual cortex parallels task performance around the visual field. eLife, 10. 10.7554/eLife.67685

Blakemore, C., Campbell, F. W. (1969). On the existence of neurones in the human visual system selectively sensitive to the orientation and size of retinal images. The Journal of Physiology, 203(1), 237–260.

Broderick, W. F., Simoncelli, E. P., Winawer, J. (2022). Mapping spatial frequency preferences across human primary visual cortex. Journal of Vision, 22(4), 3.

Burgess, A. E., Colborne, B. (1988). Visual signal detection. IV. Observer inconsistency. Journal of the Optical Society of America. A, Optics and Image Science, 5(4), 617–627.

Cannon, M. W., Jr. (1985). Perceived contrast in the fovea and periphery. Journal of the Optical Society of America. A, Optics and Image Science, 2(10), 1760–1768.

Carrasco, M., Evert, D. L., Chang, I., Katz, S. M. (1995). The eccentricity effect: target eccentricity affects performance on conjunction searches. Perception Psychophysics, 57(8), 1241–1261.

Carrasco, M., Frieder, K. S. (1997). Cortical Magnification Neutralizes the Eccentricity Effect in Visual Search. In Vision Research (Vol. 37, Issue 1, pp. 63–82). 10.1016/s0042-6989(96)00102-2

Carrasco, M., McLean, T. L., Katz, S. M., Frieder, K. S. (1998). Feature asymmetries in visual search: effects of display duration, target eccentricity, orientation and spatial frequency. Vision Research, 38(3), 347–374.

Carrasco, M., Talgar, C. P., Cameron, E. L. (2001). Characterizing visual performance fields: effects of transient covert attention, spatial frequency, eccentricity, task and set size. Spatial Vision, 15(1), 61–75.

De Valois, R. L., Albrecht, D. G., Thorell, L. G. (1982a). Spatial frequency selectivity of cells in macaque visual cortex. Vision Research, 22(5), 545–559.

De Valois, R. L., De Valois, K. K. (1988). Spatial Vision. Oxford University Press.

De Valois, R. L., Yund, E. W., Hepler, N. (1982b). The orientation and direction selectivity of cells in macaque visual cortex. Vision Research, 22(5), 531–544.

Diependaele, K., Brysbaert, M., Neri, P. (2012). How Noisy is Lexical Decision? Frontiers in Psychology, 3, 348.

Dosher, B. A., Lu, Z. L. (2000). Mechanisms of perceptual attention in precuing of location. Vision Research, 40(10-12), 1269–1292.

Duncan, R. O., Boynton, G. M. (2003). Cortical magnification within human primary visual cortex correlates with acuity thresholds. Neuron, 38(4), 659–671.

Eckstein, M. P., Pham, B. T., Shimozaki, S. S. (2004). The footprints of visual attention during search with 100% valid and 100% invalid cues. Vision Research, 44(12), 1193–1207.

Eckstein, M. P., Shimozaki, S. S., Abbey, C. K. (2002). The footprints of visual attention in the Posner cueing paradigm revealed by classification images. Journal of Vision, 2(1), 25–45.

Enroth-Cugell, C., Robson, J. G. (1966). The contrast sensitivity of retinal ganglion cells of the cat. The Journal of Physiology, 187(3), 517–552.

Falkenberg, H. K., Bex, P. J. (2007). Sources of motion-sensitivity loss in glaucoma. Investigative Ophthalmology Visual Science, 48(6), 2913–2921.

Fernández, A., Li, H.-H., Carrasco, M. (2019). How exogenous spatial attention affects visual representation. Journal of Vision, 19(10), 100b.

Fernández, A., Okun, S., Carrasco, M. (2022). Differential Effects of Endogenous and Exogenous Attention on Sensory Tuning. The Journal of Neuroscience: The Official Journal of the Society for Neuroscience, 42(7), 1316–1327.

Foster, K. H., Gaska, J. P., Nagler, M., Pollen, D. A. (1985). Spatial and temporal frequency selectivity of neurones in visual cortical areas V1 and V2 of the macaque monkey. In The Journal of Physiology (Vol. 365, Issue 1, pp. 331–363). 10.1113/jphysiol.1985.sp015776

Gold, J. M., Sekuler, A. B., Bennett, P. J. (2004). Characterizing perceptual learning with external noise. Cognitive Science, 28(2), 167–207.

Hayes, R. D., Merigan, W. H. (2007). Mechanisms of Sensitivity Loss due to Visual Cortex Lesions in Humans and Macaques. Cerebral Cortex, 17(5), 1117–1128.

Heeley, D. W., Buchanan-Smith, H. M., Cromwell, J. A., Wright, J. S. (1997). The oblique effect in orientation acuity. Vision Research, 37(2), 235–242.

Henriksson, L., Nurminen, L., Hyvärinen, A., Vanni, S. (2008). Spatial frequency tuning in human retinotopic visual areas. Journal of Vision, 8(10), 5.1–13.

Hess, R. F., & Hayes, A. (1994). The coding of spatial position by the human visual system: effects of spatial scale and retinal eccentricity. Vision Research, 34(5), 625–643.

Hilz, R., Cavonius, C. R. (1974). Functional organization of the peripheral retina: sensitivity to periodic stimuli. Vision Research, 14(12), 1333–1337.

Himmelberg, M. M., Kurzawski, J. W., Benson, N. C., Pelli, D. G., Carrasco, M., Winawer, J. (2021). Cross-dataset reproducibility of human retinotopic maps. NeuroImage, 244, 118609.

Himmelberg, M. M., Winawer, J., Carrasco, M. (2022). Linking individual differences in human primary visual cortex to contrast sensitivity around the visual field. Nature Communications, 13(1), 3309.

Hubel, D. H., Wiesel, T. N. (1962). Receptive fields, binocular interaction and functional architecture in the cat’s visual cortex. The Journal of Physiology, 160, 106–154.

Hubel, D. H., Wiesel, T. N. (1977). Ferrier lecture. Functional architecture of macaque monkey visual cortex. Proceedings of the Royal Society of London. Series B, Containing Papers of a Biological Character. Royal Society, 198(1130), 1–59.

Jakobsson, P., Lennerstrand, G. (1985). Binocular interaction in the VEP to grating stimulation. II. Spatial frequency effects. Acta Ophthalmologica, 63(3), 290–296.

Jigo, M., Tavdy, D., Himmelberg, M., Carrasco, M. (2023). Cortical magnification eliminates differences in contrast sensitivity across but not around the visual field. eLife, 12. 10.7554/eLife.84205

Kiorpes, L., Tang, C., Movshon, J. A. (1999). Factors limiting contrast sensitivity in experimentally amblyopic macaque monkeys. Vision Research, 39(25), 4152–4160.

Kleiner, M., Brainard, D., Pelli, D. (2007). What’s new in Psychtoolbox-3. Perception, 36(14), 1–16.

Klein, S. A., Levi, D. M. (2009). Stochastic model for detection of signals in noise. JOSA A, 26(11), B110–B126.

Lago, M. A., Abbey, C. K., & Eckstein, M. P. (2021). Foveated Model Observers for Visual Search in 3D Medical Images. IEEE Transactions on Medical Imaging, 40(3), 1021–1031.

Legge, G. E., Kersten, D., Burgess, A. E. (1987). Contrast discrimination in noise. Journal of the Optical Society of America. A, Optics and Image Science, 4(2), 391–404.

Levi, D. M., Klein, S. A. (1986). Sampling in spatial vision. Nature, 320(6060), 360–362.

Levi, D. M., Klein, S. A. (2002). Classification images for detection and position discrimination in the fovea and parafovea. Journal of Vision, 2(1), 46–65.

Levi, D. M., & Klein, S. A. (2003). Noise provides some new signals about the spatial vision of amblyopes. The Journal of Neuroscience: The Official Journal of the Society for Neuroscience, 23(7), 2522–2526.

Levi, D. M., Klein, S. A., Chen, I. (2008). What limits performance in the amblyopic visual system: seeing signals in noise with an amblyopic brain. Journal of Vision, 8(4), 1.1–23.

Li, H.-H., Barbot, A., Carrasco, M. (2016). Saccade Preparation Reshapes Sensory Tuning. Current Biology: CB, 26(12), 1564–1570.

Ling, S., Liu, T., Carrasco, M. (2009). How spatial and feature-based attention affect the gain and tuning of population responses. Vision Research, 49(10), 1194–1204.

Luzardo, F., Yeshurun, Y. (2021). Inter-individual variations in internal noise predict the effects of spatial attention. Cognition, 217(104888), 104888.

Lu, Z. L., Dosher, B. A. (1998). External noise distinguishes attention mechanisms. Vision Research, 38(9), 1183–1198.

Lu, Z. L., Dosher, B. A. (1999). Characterizing human perceptual inefficiencies with equivalent internal noise. Journal of the Optical Society of America. A, Optics, Image Science, and Vision, 16(3), 764–778.

Lu, Z. L., Dosher, B. A. (2008). Characterizing observers using external noise and observer models: assessing internal representations with external noise. Psychological Review, 115(1), 44–82.

Lu, Z. L., Dosher, B. A. (2023). Hierarchical Bayesian perceptual template modeling of mechanisms of spatial attention in central and peripheral cuing. Journal of Vision, 23(2), 12.

Mäkelä, P., Näsänen, R., Rovamo, J., Melmoth, D. (2001). Identification of facial images in peripheral vision. Vision Research, 41(5), 599–610.

Mareschal, I., Bex, P. J., Dakin, S. C. (2008a). Local motion processing limits fine direction discrimination in the periphery. Vision Research, 48(16), 1719–1725.

Mareschal, I., Morgan, M. J., Solomon, J. A. (2008b). Contextual effects on decision templates for parafoveal orientation identification. Vision Research, 48(27), 2689–2695.

McAnany, J. J., Park, J. C. (2018). Reduced Contrast Sensitivity is Associated With Elevated Equivalent Intrinsic Noise in Type 2 Diabetics Who Have Mild or No Retinopathy. Investigative Ophthalmology Visual Science, 59(6), 2652–2658.

Michel, M., & Geisler, W. S. (2011). Intrinsic position uncertainty explains detection and localization performance in peripheral vision. Journal of Vision, 11(1), 18.

Moraglia, G. (1989). Visual search: spatial frequency and orientation. Perceptual and Motor Skills, 69(2), 675–689.

Movshon, J. A., Thompson, I. D., Tolhurst, D. J. (1978a). Spatial and temporal contrast sensitivity of neurones in areas 17 and 18 of the cat’s visual cortex. In The Journal of Physiology (Vol. 283, Issue 1, pp. 101–120). 10.1113/jphysiol.1978.sp012490

Movshon, J. A., Thompson, I. D., Tolhurst, D. J. (1978b). Spatial summation in the receptive fields of simple cells in the cat’s striate cortex. The Journal of Physiology, 283, 53–77.

Murray, R. F., Bennett, P. J., Sekuler, A. B. (2002). Optimal methods for calculating classification images: weighted sums. Journal of Vision, 2(1), 79–104.

Nagai, M., Bennett, P. J., Sekuler, A. B. (2008). Exploration of vertical bias in perceptual completion of illusory contours: Threshold measures and response classification. Journal of Vision, 8(7), 25.1–17.

Neri, P. (2010). How inherently noisy is human sensory processing? Psychonomic Bulletin Review, 17(6), 802–808.

Neri, P., Levi, D. M. (2006). Receptive versus perceptive fields from the reversecorrelation viewpoint. Vision Research, 46(16), 2465–2474.

Ohl, S., Kuper, C., Rolfs, M. (2017). Selective enhancement of orientation tuning before saccades. Journal of Vision, 17(13), 2.

Paltoglou, A. E., Neri, P. (2012). Attentional control of sensory tuning in human visual perception. Journal of Neurophysiology, 107(5), 1260–1274.

Pelli, D. G. (1981). Effects of visual noise. http://citeseerx.ist.psu.edu/viewdoc/download?doi=10.1.1.706.822&rep=rep1&type=pdf

Pelli, D. G. (1985). Uncertainty explains many aspects of visual contrast detection and discrimination. Journal of the Optical Society of America. A, Optics and Image Science, 2(9), 1508–1532.

Pelli, D. G., Blakemore, C. (1990). The quantum efficiency of vision. Vision: Coding and Efficiency, 3–24.

Pentland, A. (1980). Maximum likelihood estimation: the best PEST. Perception Psychophysics, 28(4), 377–379.

Pointer, J. S., Hess, R. F. (1989). The contrast sensitivity gradient across the human visual field: with emphasis on the low spatial frequency range. Vision Research, 29(9), 1133–1151.

Polyak, S. L. (1941). The retina (Vol. 607). Univ. Chicago Press The retina.

Ratcliff, R., Voskuilen, C., McKoon, G. (2018). Internal and external sources of variability in perceptual decision-making. Psychological Review, 125(1), 33–46.

Rijsdijk, J. P., Kroon, J. N., van der Wildt, G. J. (1980). Contrast sensitivity as a function of position on the retina. Vision Research, 20(3), 235–241.

Robson, J. G., Graham, N. (1981). Probability summation and regional variation in contrast sensitivity across the visual field. Vision Research, 21(3), 409–418.

Rockel, A. J., Hiorns, R. W., & Powell, T. P. (1980). The basic uniformity in structure of the neocortex. Brain: A Journal of Neurology, 103(2), 221–244.

Rovamo, J., Virsu, V., Näsänen, R. (1978). Cortical magnification factor predicts the photopic contrast sensitivity of peripheral vision. Nature, 271(5640), 54–56.

Schiller, P. H., Finlay, B. L., Volman, S. F. (1976a). Quantitative studies of single-cell properties in monkey striate cortex. III. Spatial frequency. Journal of Neurophysiology, 39(6), 1334–1351.

Schiller, P. H., Finlay, B. L., Volman, S. F. (1976b). Quantitative studies of single-cell properties in monkey striate cortex. II. Orientation specificity and ocular dominance. Journal of Neurophysiology, 39(6), 1320–1333.

Shimozaki, S. S., Eckstein, M. P., Abbey, C. K. (2005). Spatial profiles of local and nonlocal effects upon contrast detection/discrimination from classification images. Journal of Vision, 5(1), 45–57.

Song, C., Schwarzkopf, D. S., Kanai, R., Rees, G. (2015). Neural population tuning links visual cortical anatomy to human visual perception. Neuron, 85(3), 641–656.

Strasburger, H., Rentschler, I., Jüttner, M. (2011). Peripheral vision and pattern recognition: a review. Journal of Vision, 11(5), 13.

Van Surdam Graham, N. (1989). Visual Pattern Analyzers. Oxford University Press.

Vilidaite, G., Baker, D. H. (2017). Individual differences in internal noise are consistent across two measurement techniques. Vision Research, 141, 30–39.

Virsu, V., Rovamo, J. (1979). Visual resolution, contrast sensitivity, and the cortical magnification factor. Experimental Brain Research. Experimentelle Hirnforschung. Experimentation Cerebrale, 37(3), 475–494.

Wardle, S. G., Bex, P. J., Cass, J., Alais, D. (2012). Stereoacuity in the periphery is limited by internal noise. Journal of Vision, 12(6), 12.

Watkins, D. W., Berkley, M. A. (1974). The orientation selectivity of single neurons in cat striate cortex. Experimental Brain Research. Experimentelle Hirnforschung. Experimentation Cerebrale, 19(4), 433–446.

Watson, A. B., & Ahumada, A. J., Jr. (2005). A standard model for foveal detection of spatial contrast. Journal of Vision, 5(9), 717–740.

Wichmann, F. A., & Hill, N. J. (2001). The psychometric function: I. Fitting, sampling, and goodness of fit. Perception & Psychophysics, 63(8), 1293–1313.

Wright, M. J., Johnston, A. (1983). Spatiotemporal contrast sensitivity and visual field locus. Vision Research, 23(10), 983–989.

Wyart, V., Nobre, A. C., Summerfield, C. (2012). Dissociable prior influences of signal probability and relevance on visual contrast sensitivity. Proceedings of the National Academy of Sciences of the United States of America, 109(9), 3593–3598.

Xu, X., Anderson, T. J., Casagrande, V. A. (2007). How do functional maps in primary visual cortex vary with eccentricity? The Journal of Comparative Neurology, 501(5), 741–755.

